# Motor deficits in the McGill-R-Thy1-APP transgenic rat model of Alzheimer’s Disease

**DOI:** 10.1101/2025.04.24.650385

**Authors:** Kyle M Roddick, Paige A Northrup, Heather M Schellinck, Richard E Brown

## Abstract

The McGill-R-Thy1-APP rat is a transgenic model of Alzheimer’s Disease (AD) which expresses APP with two mutations found in cases of familial AD, resulting in the development of amyloid pathology and cognitive deficits. Motor deficits are common symptoms of AD, emerging early in the disease, and are correlated with AD neuropathology and cognitive symptoms. This study evaluated heterozygous and homozygous McGill-R-Thy1-APP rats and their wildtype littermates for spontaneous alternation and locomotion in the T and Y mazes, and motor behaviour on an accelerating rotarod. Because rats often jumped off the rotarod, the maximum latency to fall across trials was examined. We found no genotype or sex effects in spontaneous alternation in either maze, nor a significant correlation of spontaneous alternation behaviour between the mazes. Female rats travelled greater distances than male rats in both mazes. While there was no genotype effect in the T maze on distance travelled, in the Y maze the hemizygous rats travelled shorter distances than the wildtype rats, while the homozygous rats travelled greater distances. There was a significant correlation between the distances travelled in each maze. Both heterozygous and homozygous rats performed worse than their wildtype littermates on the rotarod, while heavier rats performed worse than lighter rats, and female rats performed worse than male rats once their differences in weights were accounted for. These findings support the continued use of these rats as a model of AD and highlight the need to consider the possible confounding effect motor impairments have on other behavioural tests.

## 1.1 Introduction

Alzheimer’s Disease (AD) is an age-related neurodegenerative disorder characterized by the development of Aβ-plaques and neurofibrillary tangles (Duyckaerts et al., 2009; Lane et al., 2018; Scheltens et al., 2021). Although most AD research is focused on cognitive deficits, patients with AD also have behavioural (El Haj et al., 2017; Victoroff et al., 2018; Zvěřová, 2019), neuropsychiatric (Banning et al., 2020; Lyketsos et al., 2011), sensory (Murphy, 2019) and motor dysfunctions (Albers et al., 2015). Motor behaviour disabilities are common symptoms of AD (Buchman and Bennett, 2011) and higher levels of motor dysfunction are correlated with increased severity of cognitive symptoms (Aggarwal et al., 2006; Tian et al., 2020, 2019). Severity of motor behaviour impairments differentiates between healthy controls, patients with mild cognitive impairment, and patients with AD (Ansai et al., 2019). Walking gait (Buracchio et al., 2010; Tian et al., 2020), finger dexterity (Bologna et al., 2020; Mollica et al., 2019; Suzumura et al., 2018), and handwriting (Yu and Chang, 2019) are all impaired in AD patients, and the degree of motor impairment is associated with the level of neuropathology (Koychev et al., 2018; Nadkarni et al., 2017; Schirinzi et al., 2018). Disturbances in gait have been proposed as a method of differentiating different types of dementia (Mc Ardle et al., 2020, 2019).

Rodent models play an important role in understanding the neurobiology of AD (Do Carmo and Cuello, 2013; Drummond and Wisniewski, 2017; Götz et al., 2018; Mckean et al., 2021) and in the search for new therapeutics (Cuello et al., 2019; Puzzo et al., 2015; Snyder et al., 2016). Many transgenic rodent models have been developed which display the neuropathological hallmarks of AD, as well as the associated cognitive deficits (Pádua et al., 2024). However, it is also important to assess the neuro-behavioural, sensory, and motor functions of these models, particularly as many behavioural tests commonly used to evaluate cognitive function require the use of visual (Brown and Wong, 2007) or auditory (O’Leary et al., 2017) cues and locomotor behaviour (O’Leary et al., 2020) to perform these tests. Accordingly, motor behavioural deficits may confound the results of behavioural studies. Age related motor deficits have been shown in the APPswe/PS1dE9 (Timothy P. O’Leary et al., 2018) and 5xFAD mouse models of AD (O’Leary et al., 2020; T. P. O’Leary et al., 2018), while the 3xTg-AD mouse model has enhanced motor performance on the rotarod (Filali et al., 2012; Garvock-de Montbrun et al., 2019; Stover et al., 2015).

The McGill-R-Thy1-APP transgenic rat model of AD expresses human amyloid-β precursor protein with both the Indiana and Swedish mutations (Leon et al., 2010). Both hemizygous and homozygous transgenic rats develop intracellular accumulation of amyloid-β, with homozygous animals also developing extracellular Aβ plaques (Iulita et al., 2014; Leon et al., 2010), neuroinflammation (Hanzel et al., 2014), and dysregulation of neurotrophins (Iulita et al., 2017). Both hemizygous and homozygous rats show cognitive deficits as early as 3-months of age in the Morris Water Maze (Galeano et al., 2014; Leon et al., 2010). Changes in motor behaviour have also been observed in this rat model. Homozygous rats show motor deficits in a beam walking test, taking longer to cross the beam and making a greater number of foot slips during crossings than wildtype controls (Petrasek et al., 2018). However, no difference between homozygous rats and wildtype controls in swim speed during the Morris Water Maze test, nor in total distance traveled In the Elevated Plus Maze or the Open Field were found in this study. Other studies, while not explicitly evaluating motor behaviour, have reported no differences in swimming ability in the Morris Water Maze in either hemizygous (Galeano et al., 2018, 2014; Martino Adami et al., 2017) or homozygous McGill-R-Thy1-APP rats (Hall et al., 2018), nor differences in the total distance travelled in the open field in hemizygous (Habif et al., 2021) or homozygous rats (Orciani et al., 2023). These studies evaluated either hemizygous or homozygous rats, not both. The present study is the first to evaluate the motor behaviour of both hemizygous and homozygous McGill-R-Thy1-APP rats. Female and male transgenic rats, and their wildtype littermates, were tested in the Y and T mazes for spontaneous alternation at 12 months of age. In addition to their percent alternation, the total distance traveled was compared between genotypes. At 13 months of age they were tested on an accelerating rotarod and their latency to fall was compared. Based on the findings of impaired performance of homozygous rats on a beam walking task (Petrasek et al., 2018), we hypothesised that the homozygous rats would display motor impairments on the rotarod. There are no previous studies which explicitly examined hemizygous McGill-R-Thy1-APP rats for motor impairment, however, as previous studies using the Morris Water Maze reported no differences in swim speed (Galeano et al., 2018, 2014; Martino Adami et al., 2017), we predicted no motor deficits in the hemizygous rats.

## 2.1 Methods

### 2.2 Subjects

McGill-R-Thy1-APP transgenic rats express human amyloid-β precursor protein with both the Indiana and Swedish mutations. Expression is under control of the murine *Thy*1.2 promoter and they are on a HsdBrl:WH Wistar background (Leon et al., 2010). Hemizygous rats were obtained from the Cuello Laboratory (McGill University) and bred in our laboratory at Dalhousie University. After weaning, rats were housed with same sex littermates in groups of 2 or 3 in polyethylene cages (45cm x 24cm x 20cm) with woodchip bedding and a polyethylene tube for enrichment. Food (Purina rodent chow #5001) and water were available *ad libitum*. The colony room was under a reversed 12:12 light:dark cycle and rats were tested during the dark phase of the light dark cycle. Rats from 4 litters were tested in this experiment. The rats were 12 months of age during the T and Y maze tasks, and 13 months of age during the Rotarod task. The rats were extensively handled prior to being run in these tests. They underwent testing in a neurodevelopmental test battery from postnatal day (PND) 1 until PND 20, requiring daily handling, and were tested daily in an operant olfactometer from 3 to 7 months of age, also involving daily handling. Genotype was determined via histological examination of brain tissue for intracellular amyloid-β and amyloid-β plaques following the behavioural studies, which were done blind with respect to genotype. There were 10 wildtype (WT) rats (5 females and 5 males), 15 hemizygous (HE) transgenic rats (8 females and 7 males), and 10 homozygous (HO) transgenic rats (2 females and 8 males; see Table 1). All procedures were approved by the Dalhousie University’s University Committee on Laboratory Animals (#14-059), following the guidelines of the Canadian Council on Animal Care.

**Table 1.** Number and weights of 12 to 13 month old male and female wildtype (WT), hemizygous (+/-), and homozygous (+/+) McGill-R-Thy1-APP rats tested in spontaneous alternation and the rotarod.

| Weight (g) | Female |  |  | Male |  |  | Overall |  |  |
| --- | --- | --- | --- | --- | --- | --- | --- | --- | --- |
|  | WT | +/- | +/+ | WT | +/- | +/+ | WT | +/- | +/+ |
|  | (N=5) | (N=8) | (N=2) | (N=5) | (N=7) | (N=8) | (N=10) | (N=15) | (N=10) |
| Mean (SD) | 368<br>(18.2) | 365<br>(43.1) | 365<br>(14.8) | 625<br>(55.8) | 617<br>(61.8) | 593<br>(51.3) | 496<br>(141) | 483<br>(140) | 548<br>(107) |
| Median | 364 | 361 | 365 | 614 | 610 | 591 | 480 | 419 | 575 |
| [Min, Max] | [351,<br>393] | [308,<br>491] | [354,<br>375] | [567,<br>704] | [537,<br>683] | [526,<br>674] | [351,<br>704] | [308,<br>683] | [354,<br>674] |

### 2.3 T and Y mazes

#### 2.3.1 Apparatus

Both the T and Y mazes were constructed of transparent Plexiglas, with black paper covering on the outside of the walls. The T maze had equal-sized stem and arms (40cm length, 12cm width, 19.5cm height) without any start box. The Y maze was symmetrical with its equal sized arms (40cm length, 12cm width, 19.5cm height) forming 120° angles. The tops of the mazes were covered with transparent Plexiglas lids to prevent the rats from climbing out of the mazes. An overhead camera was used to record the rats’ behaviour, and Limelight software (Actimetrics, Lafayette, IN) was used to track the position of the rats within the mazes.

#### 2.3.2 Procedure

To test the rats for spontaneous alternation, the continuous (free-running) method was used. Animals were placed inside one of the arms of either the T or Y maze and allowed to explore for 10 minutes without any reinforcement or punishment. All of the arms were available to enter. There were no training sessions for the animals prior to the experiment; this was the first time that they were exposed to either the T or Y maze. A week after being tested in one maze, the animals were tested in the other maze using the same procedure. In order to control for the effect of familiarity due to the similarity of the mazes, half of the animals were tested in the T maze first and the other half in the Y maze first.

#### 2.3.3 Analysis

The Limelight software recorded each arm entry that the animal made during the test. An arm entry was counted when the animal’s body (the middle-point between its nose and the base of the tail) entered an arm. An alternation triplet was counted when the animal entered three different arms in succession. These triplet sets could overlap and still be counted as alternations. For example, entries of ACBABACBAB would be counted as: ACB, CBA, BAC, ACB, and CBA; therefore 5 alternations in total (Hughes, 2004). The percentage of alternations was calculated by dividing the number of alternations by the number of total entries minus 2, all multiplied by 100 (Equation 1).

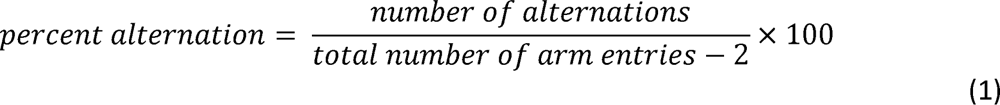

### 2.4 Rotarod

#### 2.4.1 Apparatus

Rats were tested on an AccuRotor accelerating Rotarod (Accuscan Instruments Inc. Columbus Ohio) which had an acrylic rod (7cm in diameter), separated into four 11cm sections by 11cm high Plexiglass dividers. The Rotarod accelerated from 0 to 48 rotations per minute over the course of 360s. Latency to fall was recorded by sensors in holding chambers located 36cm below the rod. The Rotarod was housed in a 1.2m x 2.6m room lit by a 60W red light.

#### 2.4.2 Procedure

Rats were tested over 5 days, receiving 6 trials each day. During each trial, rats were tested until they fell from the rod or the maximum trial length of 360s was reached. Rats were tested two at a time so that they were not in adjacent sections. Rats were weighed each day before testing.

### 2.5 Immunohistochemistry

Rats were euthanized with an overdose of sodium pentobarbital, and perfused through the left ventricle, first with saline (0.9% NaCl in dH2O, pH 7.4), followed by 4% paraformaldehyde in 0.1M phosphate buffer (pH 7.4). Brains were removed and stored in 30% sucrose in 0.1M phosphate buffer at 4°C. The brains were frozen and, using a cryostat, cut into 40µm coronal sections. The sections were then collected and stored in 0.1M phosphate buffer at 4°C.

Four sections from each rat underwent immunolabeling. Sections were permeabilized with PBS with .02% Triton X-100 (PBS-T) for 20min, followed by 20min in PBS-T with 0.3% H_2_O_2_ to quench endogenous peroxidases. After 3x 5min PBS-T washes, the sections were blocked with 30min in 5% normal goat serum, followed by overnight incubation in the primary antibody, goat anti-Ab antibody McSA1 1:500 (MédiMabs; MM-0015-P, Montreal, QC, Canada). Sections were again washed (3x 5min PBS-T) followed by the secondary antibody, biotinylated goat anti-mouse IgG 1:500 (H&L Biotin; ab6788) for 1 hour, and another wash (3x 5min PBS-T). Sections were processed with a Vectastain Elite ABC-peroxidase kit (Vector Laboratories PK-6100) for 1 hour according to manufacturer specifications, followed by another wash (3x 5min PBS-T), 15min in 0.6% 3,3’-diaminobenzidine tetrahydrochloride, 4 to 6min with 0.01% H_2_O_2_, and a final wash (3x 5min PBS-T). Sections were mounted on slides the next day, allowed to air dry, dehydrated in graded EtOH (2min each in 50%, 70%, 90%, 95%, 100%, and 100%), cleared with Histo-Clear (2x 2min), and cover slipped with Permount.

Pictomicrographs were captured using a microscope-attached digital camera system (Leica, DM E), and used to genotype the rats based on the presence of intracellular (indicating hemizygous) and extracellular (indicating homozygous) (Leon et al., 2010) Aβ reactivity (representative images are shown in S1 Fig). As genotypes were not determined until after histology was completed, experimenters were blind to the genotypes while conducting the behavioural tests.

### 2.6 Statistical Analyses

All statistical analyses were performed with the statistical program R version 4.5.0 (R Core Team, 2019). Because some rats jumped from the Rotarod into the holding chamber rather than fell, the maximum latency to fall that each rat achieved across all trials was analyzed. The mean weights of the rats during Rotarod testing were also analyzed. Linear regression models were compared using Akaike’s Information Criterion (AIC) that compares the statistical models according to their complexity and how well they fit the data (Akaike, 1974; Burnham and Anderson, 2004). Confidence intervals (95%) were calculated for all analyses.

## 3.1 Results

### 3.2 T and Y maze

#### 3.2.1 Quality checks

To ensure the quality of the data, checks were performed to confirm an unbiased assessment of spontaneous alternation (Miedel et al., 2017). A significant correlation between percent alternation and either total number of arm entries or total distance travelled could indicate that hyperdynamic locomotion had an impact on the results. Pearson correlations were run between the percent alternation, and total number of arm entries and total distance travelled, for both mazes. Percent alternation was not correlated with total number of arm entries for either the T maze (*r* = -0.07, *p* = 0.69), or the Y maze (*r* = 0.029, *p* = 0.87). Nor was percent alternation correlated with total distance travelled for either the T maze (*r* = -0.042, *p* = 0.81), or the Y maze (*r* = 0.066, *p* = 0.7). Therefore, there is no indication of hyperdynamic locomotion impacting the results.

ANOVAs were run on both mazes to examine if any of the arms were entered more times than the others, an indicator of potential environmental cues in the maze leading to biased behaviour (Miedel et al., 2017). There were no significant differences in the number of times each arm was entered for either the T maze (*F*_(2,102)_ = 0.745, *p* = 0.477, *η_G_*^2^ = 0.014) or the Y maze (*F*_(2,102)_ = 0.28, *p* = 0.756, *η_G_*^2^ = 0.005). Therefore, there is no indication of environmental cues biasing the behaviour of the rats.

#### 3.2.2 Spontaneous alternation

Linear regression models of percent alternation in the T and Y mazes were compared using AIC to select the best model for each maze. For the T maze, the null model best fit the data (S1 Table) indicating no significant effect of genotype or sex (Fig 1A).

**Fig 1.**
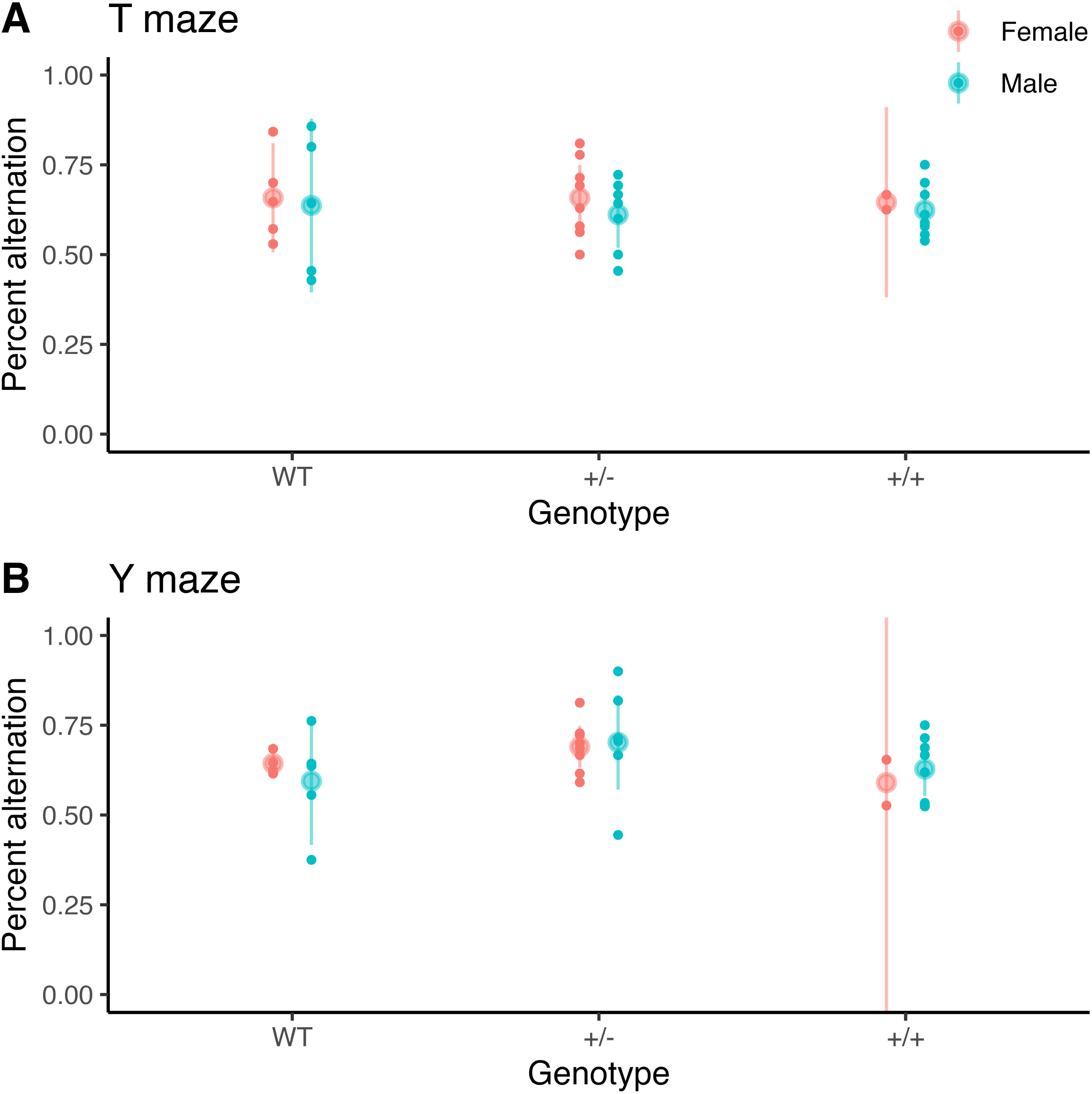
Percent Alternation in the T and Y Mazes. Mean (± CI) percent alternation in the (A) T maze and the (B) Y maze. There were no significant effects of genotype nor sex on percent alternation in either maze.

For the Y maze, the model with an effect of genotype best fit the model, however, this model did not differ significantly from the null model (*F* = 2.5, *p* = 0.096; S2 Table) again indicating no effects of genotype or sex (Fig 1B).

A Pearson correlation coefficient between the percent alternation by all the rats in the two mazes was not significant (*r* = -0.24, *p* = 0.17; S2 Fig A).

#### 3.2.3 Distance travelled

Linear regression models of the distances travelled in the T and Y mazes were compared using AIC to select the best model for each maze. For the T maze the model with just an effect of sex was the best model (S3 Table), and this model differed significantly from the null model (*F* = 11.00, *p* = 0.002). This model indicates that the male rats travelled less distance (33.8 ± 8.11 m) than the female rats (43.6 ± 9.18 m) in the T maze (CI^95^ of the difference: -15.8 - -3.9 m; Fig 2A).

**Fig 2.**
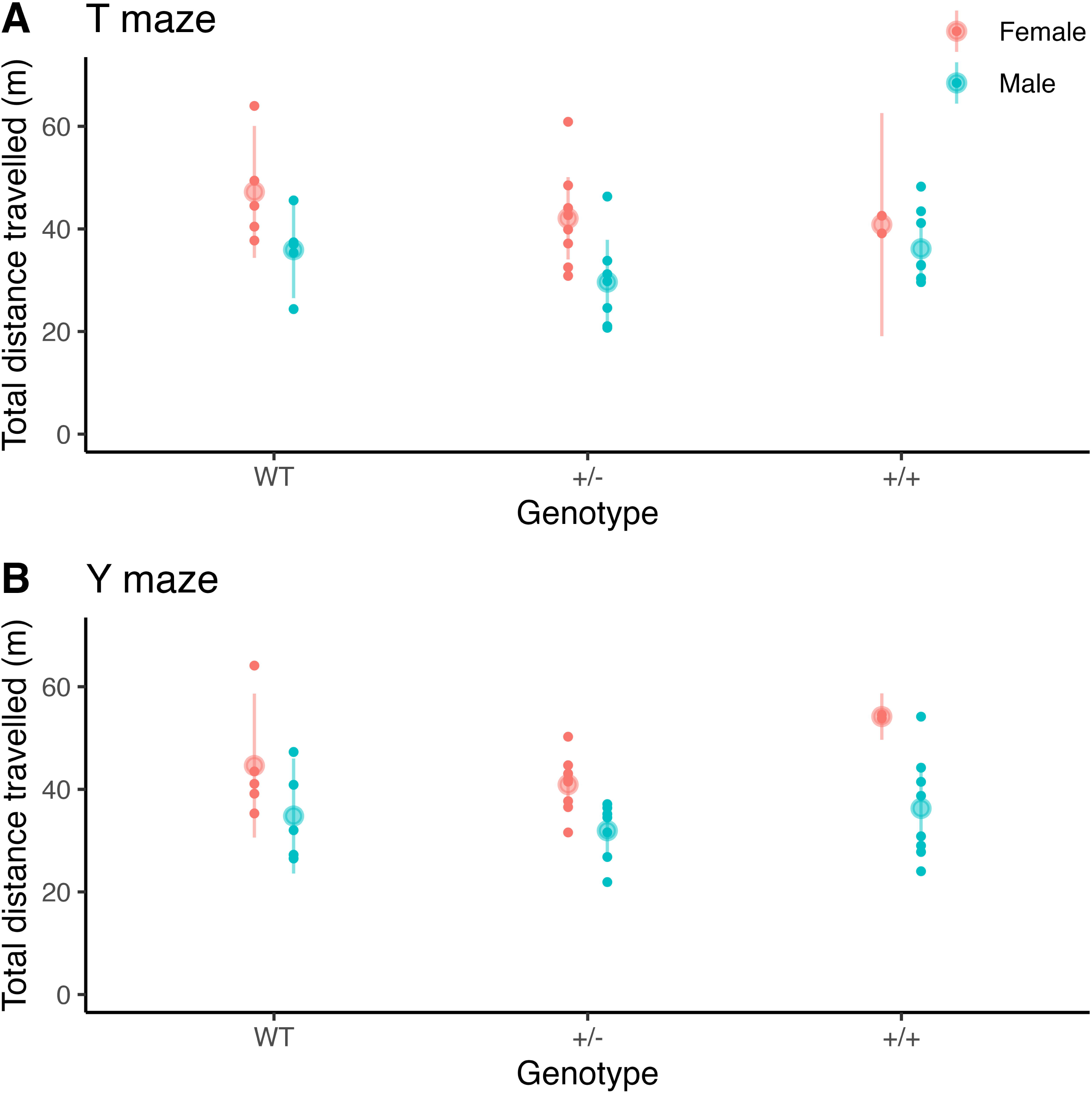
Distance Travelled in the T and Y Mazes. Mean (± CI) total distance travelled in (A) the T maze and (B) the Y maze. The female rats traveled greater distances than the male rats in both mazes. In the Y maze, the hemizygous (+/-) rats travelled less than the wildtype (WT) rats, and the homozygous (+/+) rats travelled more than the wildtype rats.

For the Y maze the model with effects of genotype and sex was the best model (S4 Table), and this model differed significantly from the null model (*F* = 5.2, *p* = 0.005). Again, the male rats (34.4 ± 8.3 m) travelled less distance than the female rats (43.9 ± 8.54 m; CI^95^ of the difference: -17 - -5.1 m; Fig 2B). This model also indicates that the hemizygous (+/-) rats travelled less than the wildtype (WT) rats (CI^95^ of the difference: -10.2 - 3.4), while the homozygous (+/+) rats travelled further than the WT rats (CI^95^ of the difference: -4.18 - 11).

### 3.3 Rotarod

The time to fall from the rotarod for each rat on each of six trials per day for five days, with LOESS (locally estimated scatterplot smoothing); (Cleveland, 1979) trend lines and 95% confidence intervals (CI^95^) for each genotype, is shown in Fig 3. Early in the study, the rats began to jump from the rotarod, resulting in frequent very low latencies and decreasing latencies to fall across trials. With the exception of one hemizygous female that achieved perfect performance by the final day of testing (S4 Fig A) rats failed to show increases in latency to fall across days as would be expected for motor learning on the rotarod (T. P. O’Leary et al., 2018). In order to best evaluate locomotor ability, the effects of sex, genotype, and body weight on the maximum latency to fall that each rat achieved across all trials were analyzed.

**Fig 3.**
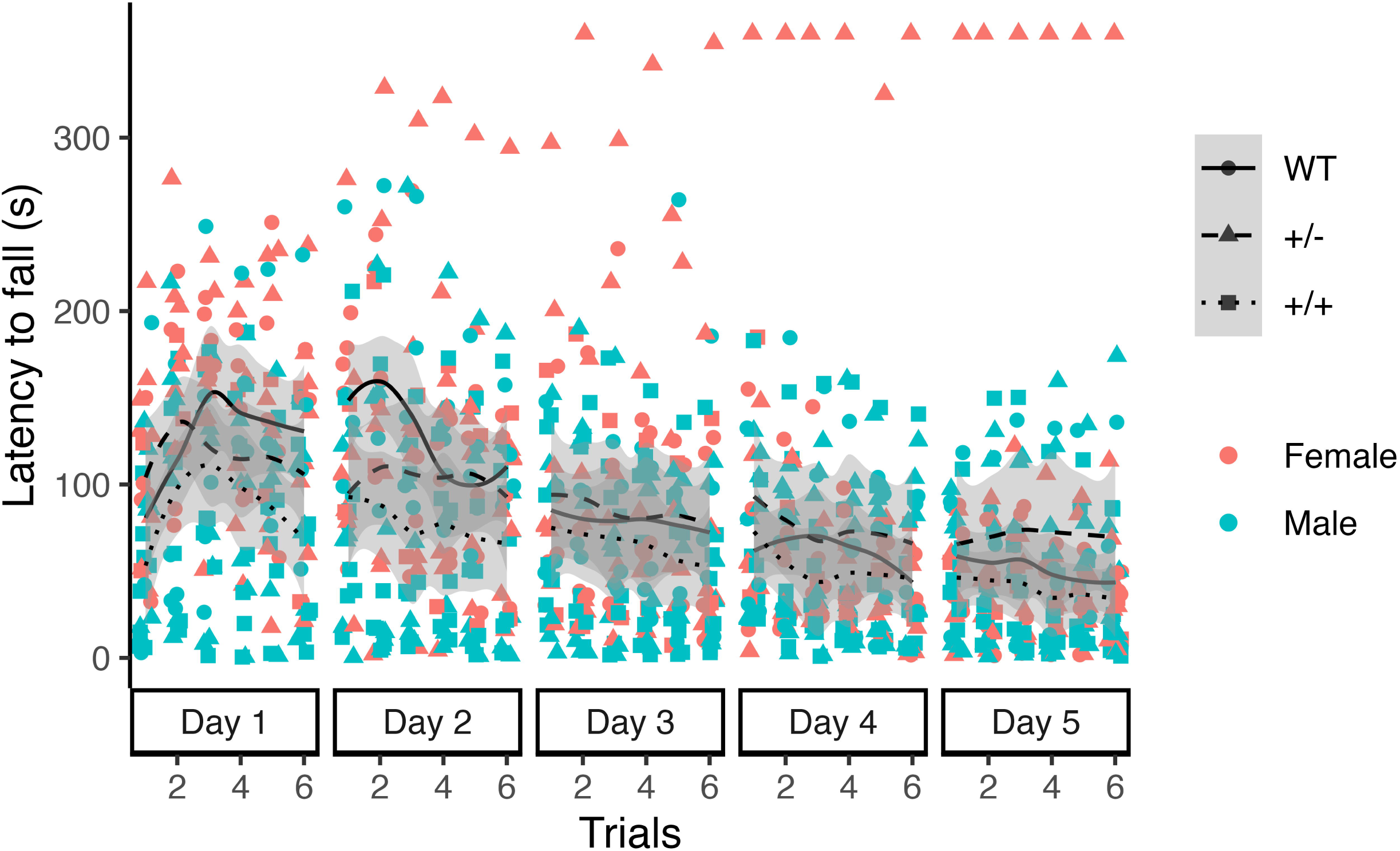
Latency to Fall from the Rotarod. Latency(s) to fall from the Rotarod across the six trials on each of five days of testing. Points for each individual trial and loess lines, with CI^95^ for each genotype, are shown. Latencies for all genotypes began to fall in the second half of the first day of testing, and continued to decrease each day. This was because the rats jumped off the Rotarod rather than fell off.

Linear regression models of the mean weights of the rats (see Table 1) across testing days were compared using AIC (S5 Table). The best model of the data (AIC = 372.4, weight = 0.75) included no effect of genotype on body weight, but did include an effect of sex with males weighing more than females (CI^95^ 209 – 274 g). This model differed significantly from the null model (*F* = 228, *p* < 0.0001). Linear regression models of the maximum latency to fall for each rat were compared using AIC (S6 Table). The additive model with effects of mean body weight, sex, and genotype was found to best model the data (AIC = 378.13, weight = 0.286), and differed significantly from the null model (*F* = 8.3, *p* < 0.001). This model indicated that body weight was negatively associated with the maximum latency to fall with lighter rats having longer latencies to fall than heavier rats (CI^95^ -1.091 – -0.332 s/g) This means that the model predicts a decrease in the latency to fall of approximately .7 seconds for each gram of body weight (s/g). The model also shows males having longer latencies to fall than females (CI^95^ 13.71 – 214.09 s), and both hemizygous (CI^95^ -69.90 – 11.61 s) and homozygous rats (CI^95^ -98.04 – -4.70 s) having shorter latencies to fall than wildtype rats. S3 Fig shows raincloud plots for the genotype differences in maximum latency to fall over the 30 trials, while Fig 4 shows that the lighter rats stayed on the rotarod longer than heavier rats for both females (*r* = -0.60, *p* = 0.017) and males (*r* = -0.49, *p* = 0.028).

**Fig 4.**
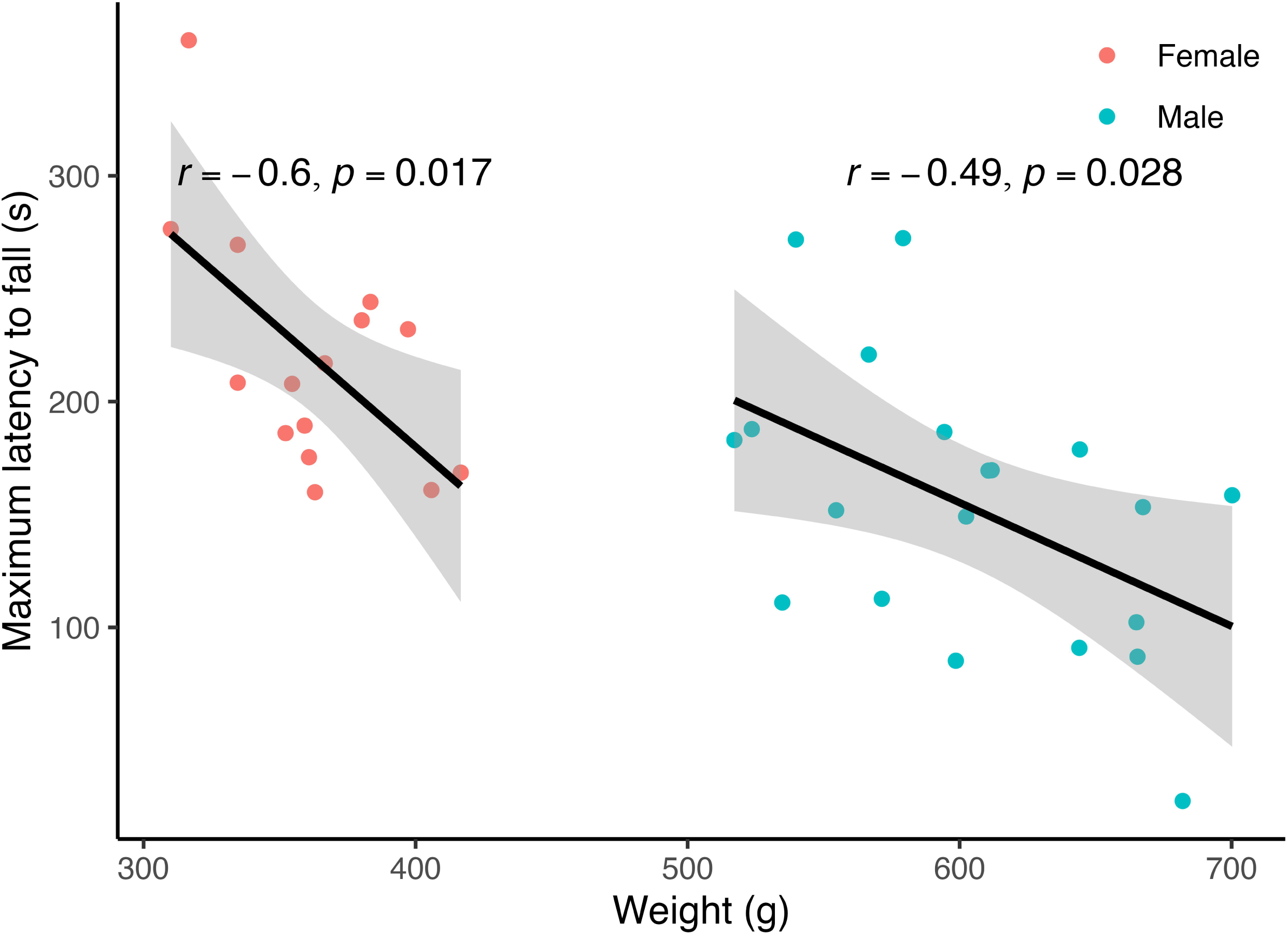
Maximum Latency to Fall from the Rotarod. Maximum latency to fall (s) from the Rotarod for individual female and male rats according to their body weight (g). Linear regression lines (± CI^95^) of the effect of weight on maximum latency to fall for females and males are shown separately because of the sex differences in body weight.

### 3.4 Cross test correlations

Pairwise Pearson correlations were run between the three measures of locomotor activity, distance travelled in the T and Y mazes and the maximum latency to fall from the Rotarod. While the distance travelled in the T and Y mazes were positively correlated (*r* = 0.65, *p* < 0.0001), the maximum latency to fall from the Rotarod was not significantly correlated with distance travelled in either the T (*r* = 0.3, *p* = 0.08) or Y maze (*r* = 0.061, *p* = 0.73; Fig 5).

**Fig 5.**
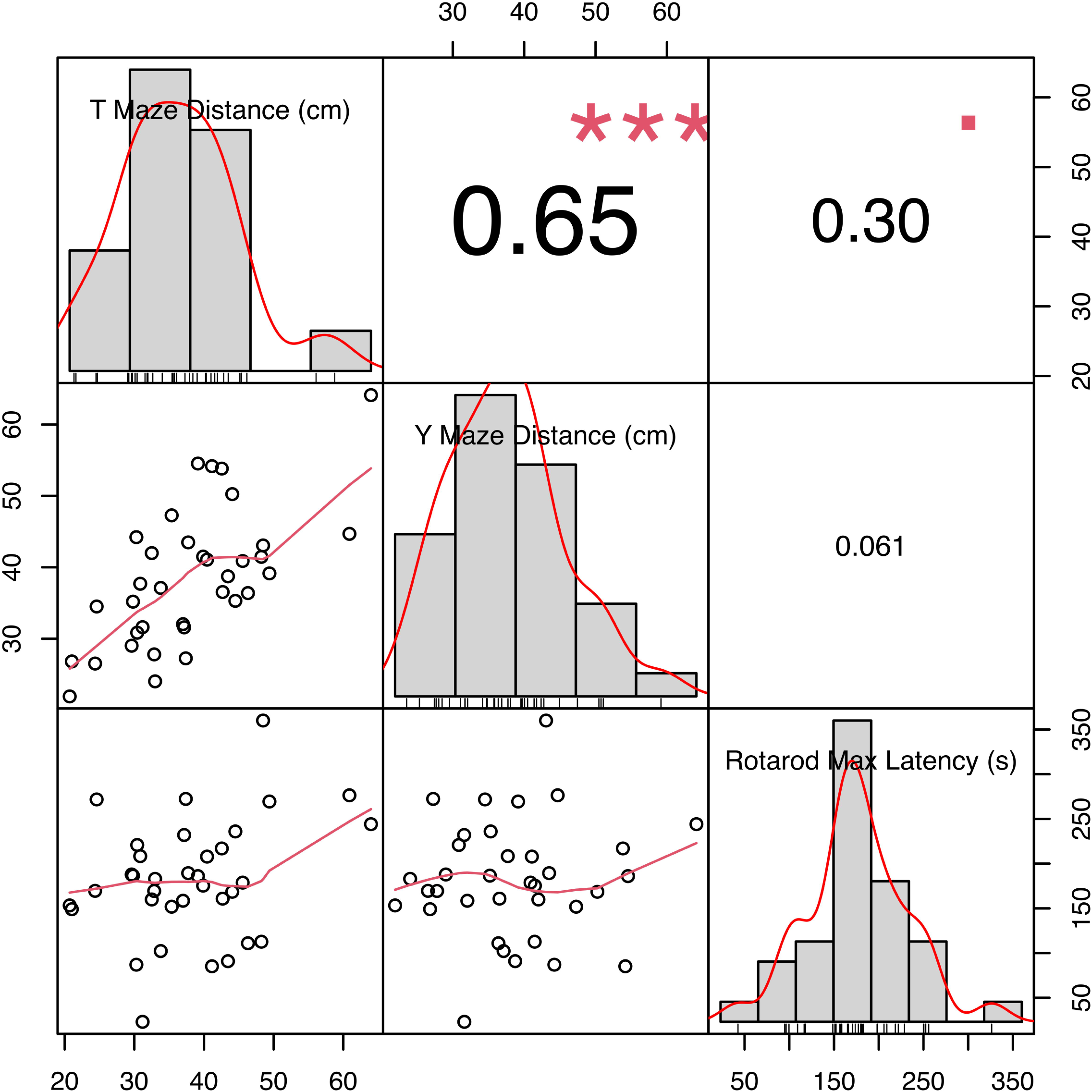
Cross Test Correlations. Pearson correlation matrix of locomotor measures. The plots in the central diagonal show histograms of the data for each variable overlayed with density curves. The plots in the lower half scatterplots of the two variables with loesses lines of best fit, while the numbers in the upper half indicate the correlation coefficient with significance level (square for p < 0.1, *** for p < 0.001).

## 4.1 Discussion

This study found no evidence of differences in either the homozygous or hemizygous McGill-R-Thy1-APP transgenic rats on spontaneous alternation compared to wildtype rats. This is in contrast to Galeano et al. (2014), who found spontaneous alternation deficits in hemizygous McGill-R-Thy1-APP rats at 6 and 12 months of age, but not at 3 months. Other rat models of AD have been tested for spontaneous alternation with mixed results. TgF344-AD transgenic rats, with human APP containing the Swedish mutation and Δ exon 9 mutant human presenilin-1 (Cohen et al., 2013), showed impaired spontaneous alternation at 18 months of age (Yang et al., 2022). Aβ-injected rats had lower rates of spontaneous alternation than controls in some studies (Bagheri et al., 2011; Cioanca et al., 2013; Hritcu et al., 2014; Wang et al., 2019; Yin et al., 2010), but not in others (Sipos et al., 2007). However, aluminum chloride induced AD model rats (Olajide et al., 2017; Zaher et al., 2020) and a streptozotocin induced AD rat model (Shi et al., 2017) had lower rates of spontaneous alternation than controls.

The present study found no significant correlation between spontaneous alternation in the T and Y mazes, but there was a significant, positive correlation between distance traveled in the two mazes. In both mazes the male rats travelled shorter distances than the female rats, but there was no clear genotype pattern between the mazes. Thus, there is little evidence of a motor impairment in either the hemizygous or homozygous McGill-R-Thy1-APP rats in the T or Y mazes.

However, there was clear evidence that homozygous McGill-R-Thy1-APP rats showed impaired motor behaviour on the rotarod compared to their wildtype littermates. The evidence also strongly suggested an impairment in motor behaviour on the rotarod in the hemizygous transgenic rats. The finding of impaired motor behaviour in the homozygous rats is in agreement with the results of Petrasek et al. (2018), who found homozygous rats to be impaired on a beam walking task. While some previous studies have tested hemizygous rats in the Morris Water Maze and reported no differences in swimming speed (Galeano et al., 2018, 2014; Martino Adami et al., 2017), no prior studies have explicitly evaluated the motor behaviour of hemizygous rats. It has been argued that the hemizygous McGill-R-Thy1-APP rats are a good model of the early stages of AD (Galeano et al., 2014) and this argument is supported by the present evidence of motor impairments on the Rotarod in the hemizygous rats. As motor dysfunction is a well established symptom of early AD (Buchman and Bennett, 2011), animal models of early AD should also be expected to display motor dysfunction.

We analyzed the single best trial for each rat, a measure which has been previously used to assess motor behaviour (Allard et al., 2018), because the rats frequently jumped off of the rod during testing rather than falling off. The data support this observation, with only one rat showing a clear improvement over the test (S4 Fig A). While there were 104 trials with a latency of less than 10 seconds to fall, including 23 trials with a latency of less than 2 seconds. All three genotypes began to show a decline in performance during the first day of testing (Fig 3), indicating that jumping from the rotarod started to occur very early during testing. This jumping behaviour means that in many cases the latency to fall is not an accurate measure of how long the rat was able to stay on the rod. By taking the longest latency for each individual rat we were able to get a better measure of the capacity of the rat to stay on the rod. While 54% of the rats had their maximum latency trial on the first day of testing, no rats had their maximum latency on the first trial. It is possible that the advanced age of the rats when tested (13 months) made the repeated trials of the task more tiring for them, and that this lead to the jumping behaviour. Other studies have reported instances of passive rotation on the Rotarod (Hamm et al., 1994; Meconi et al., 2018), where the rats grip the rod and rotate with it, but no instances of passive rotation were observed in the present study. We found no other published data on rats jumping from the rotarod, though it has been observed in mice (Schönfeld et al., 2017), so there are no established methods for dealing with this type of animal “misbehaviour” (Breland and Breland, 1961).

The method of analyzing the maximum latency to fall for each rat is not without fault. The worst performing rat, a hemizygous male, had a maximum latency to fall of just 23.1 sec, and three trials with a latency less than 1 second (S4 Fig B). While this rat was the second heaviest rat tested, it is unlikely that the rotarod was able to measure the true motor capacity of this animal. However, we found a clear effect of weight in this study, with lighter rats performing better on the rotarod than heavier rats, which is in agreement with what we have found when testing mouse models of AD (Garvock-de Montbrun et al., 2019; O’Leary et al., 2020; Timothy P. O’Leary et al., 2018; T. P. O’Leary et al., 2018; Stover et al., 2015). This suggests that using the maximum latency each rat achieved is accurately measuring the effects of genotype and sex on of motor function.

When corrected for body weight, the model of the latencies to fall indicated that males stayed on the rod longer than females. This is in contrast with the uncorrected results of the experiment which found that female rats had a greater maximum latency to fall (219 ± 54 s) than males (153 ± 62 s). This apparent contradiction is due to the combination of the effect of weight on the latency to fall, and the difference in weights between female and male rats. Our model found that the greater the weight of the rat, the shorter the time it was able to stay on the rod. We also found a substantial difference in the weights of female and male rats (Table 1, Fig 4). There was no overlap in weights between the two sexes, with the heaviest female (417g) weighing 100g less than the lightest male (517g). So while the males did not stay on the rod as long as the females, they performed better than the females once their weight was taken into account. Studies of sex differences in rotarod performance in rat and mouse models of AD found that female rats performed better than male rats (Hernandez et al., 2020), though the effect of body weight was not accounted for in that study. When rats were matched by weight, female rats also performed better than males (O’Connor et al., 2003). We previously found no effect of sex on rotarod performance in 3xTg-AD mice at either 6 (Stover et al., 2015), or 16 months of age (Garvock-de Montbrun et al., 2019) once weight was taken into account. Female 5xFAD mice performed better than males (O’Leary et al., 2020), however this study did not account for differences in their body weight in the analysis of sex differences.

Other rat models of AD have been tested on the rotarod with mixed results. Some studies using a model of AD induced by Aβ injection found no deficits in latency to fall from the Rotarod (Huang et al., 2011; Navabi et al., 2018), while other studies using an Aβ-induced model did find deficits in Rotarod performance (Kuo et al., 2021; Lee et al., 2014; Prakash et al., 2013). Deficits in motor performance on the rotarod have also been found in models of AD induced by aluminum chloride exposure (Justin-Thenmozhi et al., 2018; Lakshmi et al., 2015), trimethyltin chloride (Hosseini et al., 2020), and scopolamine (Manral et al., 2016). Meanwhile, a study using a streptozotocin induced AD model (Stanojevic et al., 2022), and a study using the App^NL-G-F^ transgenic rat (Pang et al., 2022), with three familiar App mutations (Swedish, Beyreuther/Iberian, and Arctic) and a humanized Aβ sequence knocked into the rat App gene, found no differences on the rotarod at five months of age. These findings highlight the need for studies to consider the possible impact motor deficits may have when performing cognitive tests on AD models. Many tests, such as the Morris Water Maze, the T maze, and the Y maze, require the animal to move around the testing area. If they have motor impairments, a lower performance could be due to the motor impairments rather than a cognitive deficit. Conversely, if an experimental intervention restores cognitive function, but is not effective for motor function, these tests may falsely find a negative result if motor function impacts the animal’s ability to complete the test.

Different genotype effects were found in distance travelled in the T and Y mazes versus the maximum latency to fall from the rotarod. There was also no significant correlation between the maximum latency to fall from the rotarod and the distance travelled in either maze. This is likely due to the distance travelled in the T and Y mazes being a measure of voluntary movement, while the maximum latency to fall from the rotarod is a measure of forced movement. Lambert et al. (1996) found that performance of rats on a forced movement treadmill task did not predict their voluntary movement in a home-cage running wheel. Additionally, a study examining the motor performance of seven inbred strains of mice found no correlation between the strains performance on a forced treadmill task and voluntary wheel running (Lerman et al., 2002).

Although the rotarod is a commonly used apparatus for measuring motor learning and coordination in mice and rats (Schönfeld et al., 2017), there are a number of parameters which can affect the performance of rodents on the rotarod. The use of an accelerating vs fixed speed rotarod, and their rate of acceleration or speed, respectively, will influence the sensitivity of the test (Nathan R. Rustay et al., 2003). The diameter of the rod (if it is small enough for the rodent to grab hold of and passively rotate) can also effect performance (Nathan R Rustay et al., 2003). Differences in performance can also be seen between rotarods from different manufacturers (Dreesen and Riedel, 2018), which could be due to differences in the rate of acceleration of the rod (Bohlen et al., 2009). The training schedule can also effect performance, with rodents showing a learning curve over trials (Garvock-de Montbrun et al., 2019; O’Leary et al., 2020; Nathan R Rustay et al., 2003).

Few experiments test aged rodents on the rotarod, and since AD is a disease of aging, it is important to examine the age-related decline in motor behaviour (Garvock-de Montbrun et al., 2019; O’Leary et al., 2020). Although mice gain weight with age, the increase is from 26-34 g at 6 months of age (Stover et al., 2015) up to 26-46 g at 16 months of age (Garvock-de Montbrun et al., 2019). This is a very small age-related increase in body weight compared rats, which increase from 179-349 g in young rats (Cartmell et al., 1991) to the 308-704 g found in the present study. As with mice (Stover et al., 2015), heavier rats fell faster from the rotarod than lighter rats (Rozas et al., 1997).

Two behaviours which have resulted in animals being excluded from rotarod studies involve passive rotations and failure to learn over trials. Rodents can cling to the rod and show passive rotations (Bohlen et al., 2009; Deacon, 2013) and Shan et al (2023) excluded 10 of 58 C57BL/6 mice due to “bad performance in baseline testing” on the rotarod. Zieglowski et al. (2020) excluded 14 of 30 Wistar Han rats due to “lack of learning ability” and Cartmell et al. (1991) excluded 25 of 60 Sprague Dawley rats for failing to show improved over 10 trials on the rotarod. These problems came as a surprise to us as we had not had mice show passive rotations nor have we had to remove mice for failing to learn over repeated trials on the rotarod (Garvock-de Montbrun et al., 2019; O’Leary et al., 2020; Stover et al., 2015). We have even recommended the rotarod as one of the most reliable tests for measuring motor function in mice (O’Leary et al., 2020). The standard rotarod is not designed for these large rats, which easily jump down from the rod without waiting to fall. Although we modified the standard procedure by testing only two rats at a time, they quickly learned to jump off the rod and other modifications are necessary (Bohlen et al., 2009; Cartmell et al., 1991; Schönfeld et al., 2017). Some rotarod systems are capable of delivering a mild electrical shock to the rodents when they fall off the rod and land in the holding chambers below (Rozas et al., 1997), it is possible that such a system would discourage such jumping behaviour, leading to a more typical increase in latency to fall across trials and more reliable measure of age-related motor dysfunction.

The findings of this study, particularly with regards to the effects of sex in the homozygous rats, should be interpreted with some caution due to the combination of the jumping off behaviour and the imbalanced sex ratio of the homozygous rats (2 females and 8 males). The jumping off behaviour prevents the examination of motor learning which is typically assessed in studies using the rotarod, and while the use of the maximum latency to fall as the main dependent measure appears to have been an effective method of assessing the motor ability of these rats, as shown by the detection of weight effects similar to those found in other studies, it remains a cruder measure. The analysis of the maximum latency to fall from the rotarod found main effects of genotype and sex, but no significant interaction between these two factors. However, due to the imbalanced sex ratio among the homozygous rats this lack of an interaction should be interpreted with particular caution.

In summary, this study found evidence of motor deficits in both the homozygous and hemizygous transgenic McGill-R-Thy1-APP rat models of AD. While Petrasek et al. (2018) also found impairment in the homozygous rats, this is the first study to find such deficits in the hemizygous rats. These findings support the use of the McGill-R-Thy1-APP as a model of AD, though it is important to consider the potential effects of motor impairments when assessing other behaviours in this, or any other rodent AD model.

## Data Availability

The data set and analysis scripts used in this study are available on the Mount Allison University Dataverse on Borealis at https://doi.org/10.5683/SP3/8HHKPA.

## CRediT Author contributions

KMR: Conceptualization; Data curation; Formal analysis; Investigation; Methodology; Visualization; Writing - original draft; Writing - review & editing. PAN: Investigation; Methodology; Writing – review & editing. HMS: Conceptualization; Project administration; Methodology; Resources; Supervision; Writing - review & editing. REB: Conceptualization; Funding acquisition; Project administration; Resources; Supervision; Writing - review & editing.

## Acknowledgments

Thanks to Amirreza Shahisavandi for assistance with behavioural testing.

