## Supplementary material for "Motor deficits in the McGill-R-Thy1-APP transgenic rat model of Alzheimer’s Disease": S1 Fig

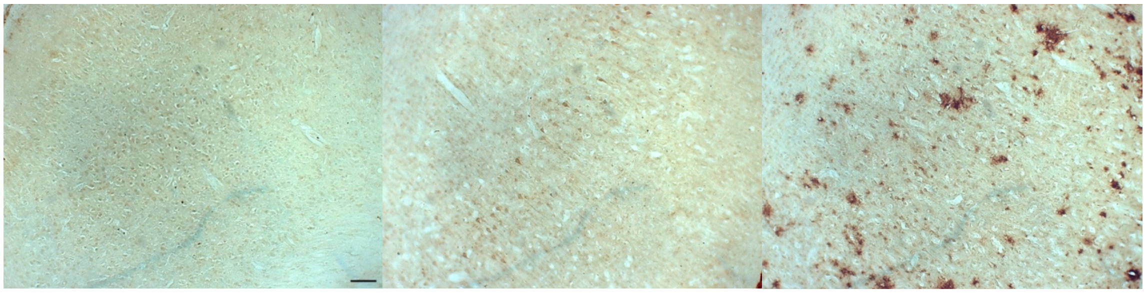

**S1 Fig. Representative images of Aβ pathology.** Representative pictomicrographs showing Aβ immunoreactivity (brown) in wildtype (left), hemizygous (center), and homozygous (right) rat cortex. Scale bar: 100μm.
