## Supplementary material for "Motor deficits in the McGill-R-Thy1-APP transgenic rat model of Alzheimer’s Disease": S1 Table

**S1 Table. Model selection table for percentage of alternations in the T maze.** Akaike's Information Criterion (AIC) model comparison of percentage of alternations in the T maze. Models are ranked from lowest to highest AIC values, and the degrees of freedom (df) used by the model calculated. The delta value indicates the difference between the AIC of a model and the AIC of the model with the lowest AIC. The weight of the models indicates the probability of the particular model out of all tested models.

| geno | sex | geno:sex | df | logLik | AIC | delta | weight |
|---|---|---|---|---|---|---|---|
| | | | 2 | 28.32 | -52.64 | 0.00 | 0.555 |
| | + | | 3 | 28.75 | -51.50 | 1.14 | 0.314 |
| + | | | 4 | 28.40 | -48.80 | 3.84 | 0.081 |
| + | + | | 5 | 28.79 | -47.58 | 5.06 | 0.044 |
| + | + | + | 7 | 28.85 | -43.69 | 8.95 | 0.006 |
