## Supplementary material for "Motor deficits in the McGill-R-Thy1-APP transgenic rat model of Alzheimer’s Disease": S2 Fig

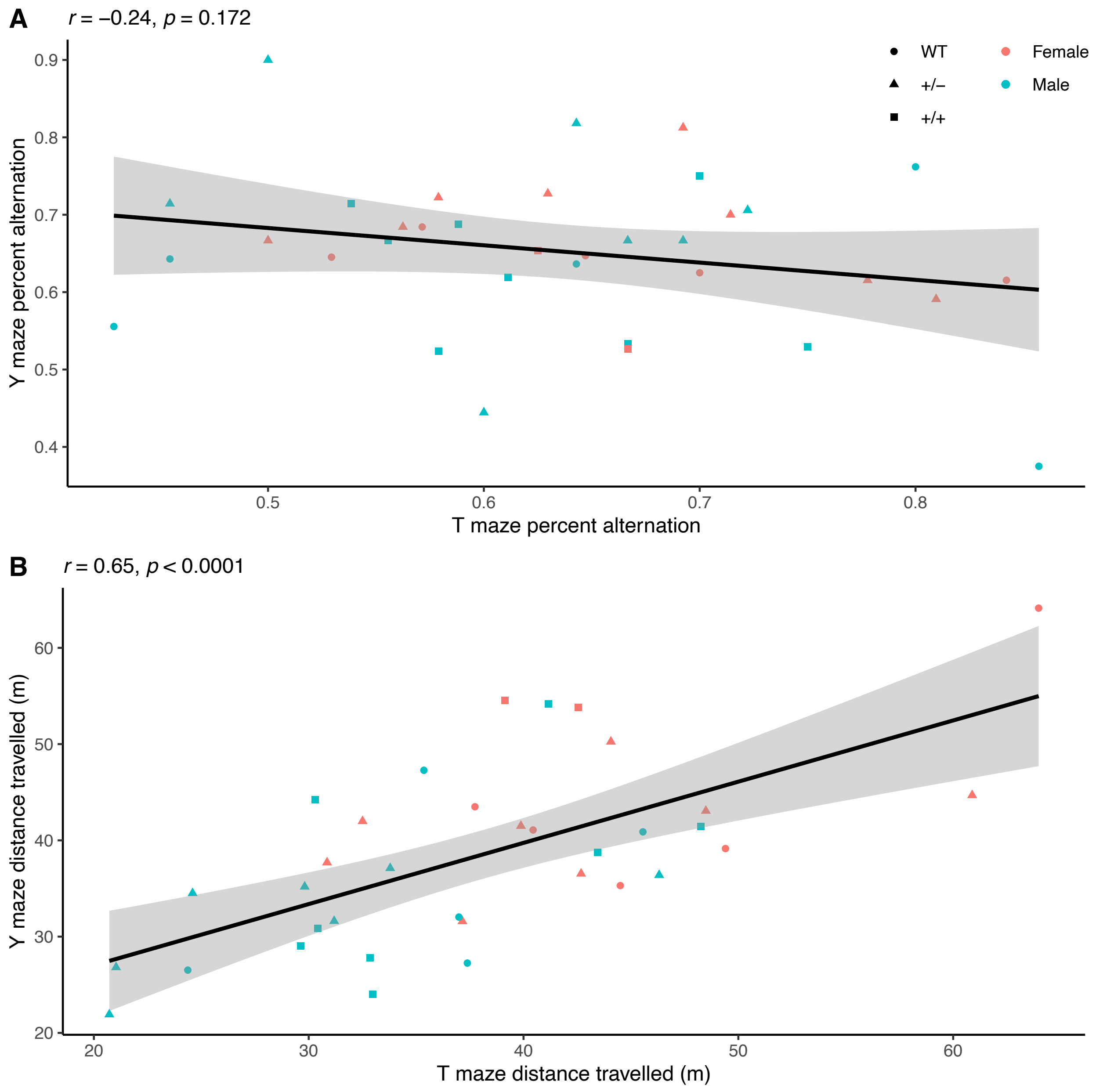


**S2 Fig. Correlations Between the T and Y Mazes.** Pearson correlation between (A) the percent alternation, and (B) the total distance travelled, by each rat in the T and Y mazes. Wildtype rats (WT) are indicated with circles, hemizygous rats (+/-) with triangles, and homozygous rats (+/+) with squares. Female rats are shown in red, while male rats are shown in blue. There was not a significant correlation between percent alternation in the two mazes, but there was a significant positive correlation between the distances travelled in the two mazes.
