## Supplementary material for "Motor deficits in the McGill-R-Thy1-APP transgenic rat model of Alzheimer’s Disease": S2 Table

**S2 Table. Model selection table for percentage of alternations in the Y maze**. Akaike's Information Criterion (AIC) model comparison of percentage of alternations in the Y maze. Models are ranked from lowest to highest AIC values, and the degrees of freedom (df) used by the model calculated. The delta value indicates the difference between the AIC of a model and the AIC of the model with the lowest AIC. The weight of the models indicates the probability of the particular model out of all tested models.

| geno | sex | geno:sex | df | logLik | AIC | delta | weight |
|---|---|---|---|---|---|---|---|
| + | | | 4 | 32.84 | -57.67 | 0.00 | 0.44 |
| | | | 2 | 30.27 | -56.54 | 1.13 | 0.25 |
| + | + | | 5 | 32.84 | -55.68 | 2.00 | 0.16 |
| | + | | 3 | 30.37 | -54.74 | 2.94 | 0.10 |
| + | + | + | 7 | 33.34 | -52.68 | 4.99 | 0.04 |
