## Supplementary material for "Motor deficits in the McGill-R-Thy1-APP transgenic rat model of Alzheimer’s Disease": S3 Fig

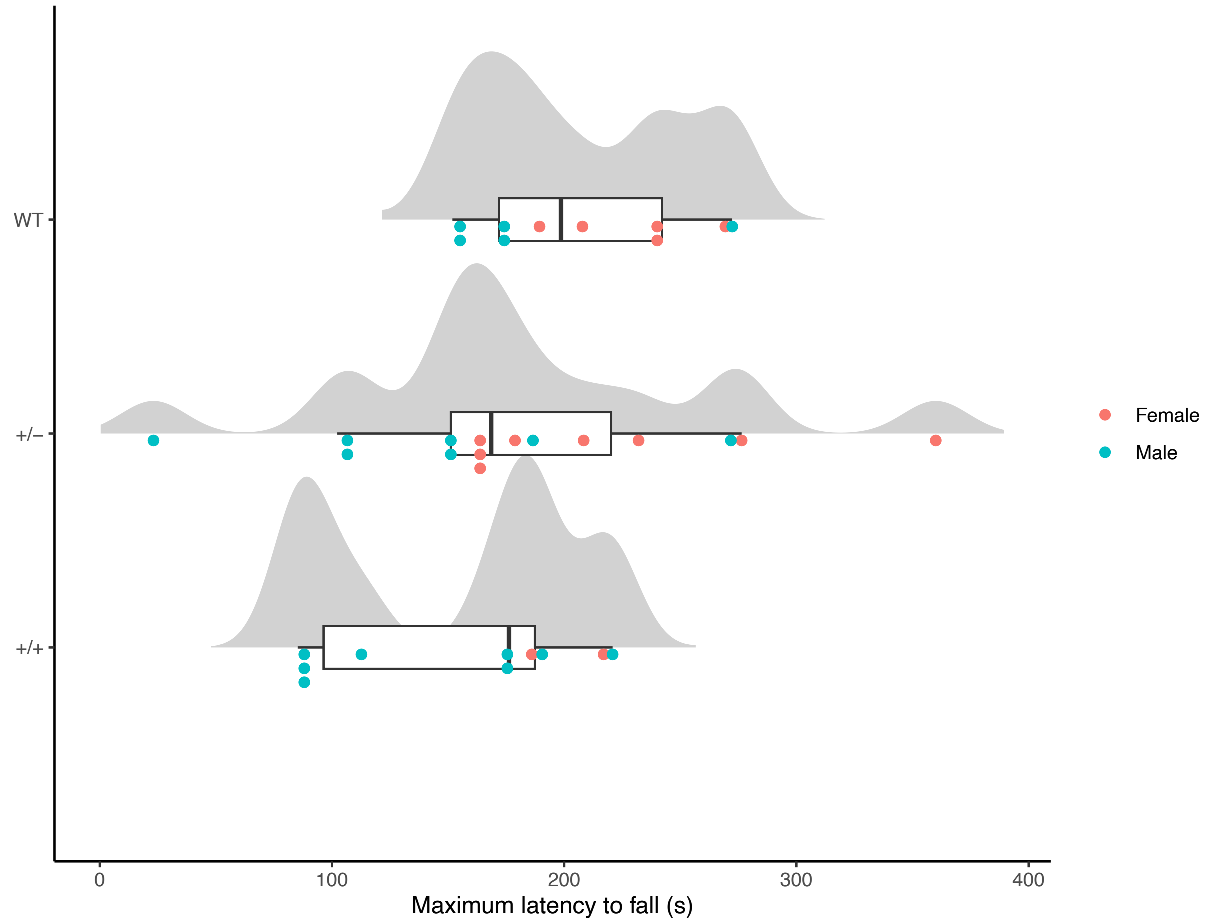


**S3 Fig. Distribution of the Maximum Latency to Fall from the Rotarod.** Raincloud plots of the maximum latency to fall (s) from the Rotarod over 30 trials for wildtype (WT), hemizygous (+/-), and homozygous (+/+) rats, showing the distribution of the data with density curves, boxplots, and data points.
