## Supplementary material for "Motor deficits in the McGill-R-Thy1-APP transgenic rat model of Alzheimer’s Disease": S3 Table

**S3 Table. Model selection table for distance travelled in the T maze.** Akaike's Information Criterion (AIC) model comparison of distance travelled in the T maze. Models are ranked from lowest to highest AIC values, and the degrees of freedom (df) used by the model calculated. The delta value indicates the difference between the AIC of a model and the AIC of the model with the lowest AIC. The weight of the models indicates the probability of the particular model out of all tested models.

| geno | sex | geno:sex | df | logLik | AIC | delta | weight |
|---|---|---|---|---|---|---|---|
| | + | | 3 | -123.86 | 253.72 | 0.00 | 0.530 |
| + | + | | 5 | -122.22 | 254.43 | 0.71 | 0.371 |
| + | + | + | 7 | -121.66 | 257.33 | 3.60 | 0.087 |
| | | | 2 | -128.98 | 261.96 | 8.24 | 0.009 |
| + | | | 4 | -127.98 | 263.95 | 10.23 | 0.003 |
