## Supplementary material for "Motor deficits in the McGill-R-Thy1-APP transgenic rat model of Alzheimer’s Disease": S4 Fig

A

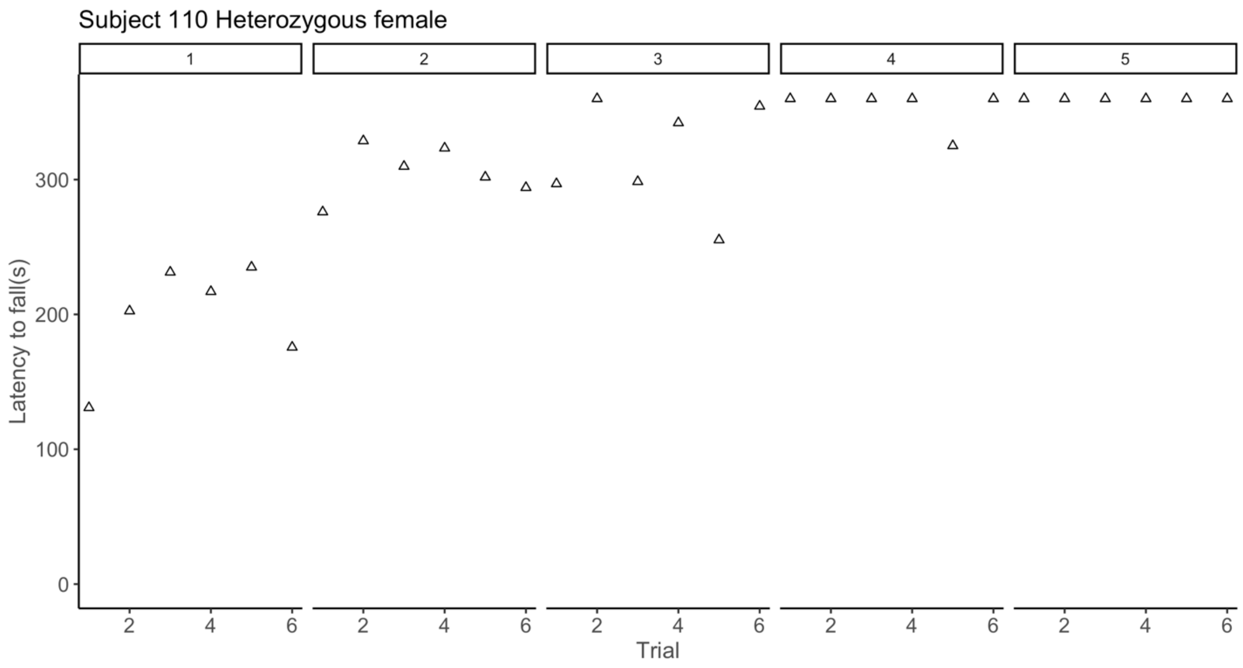


B

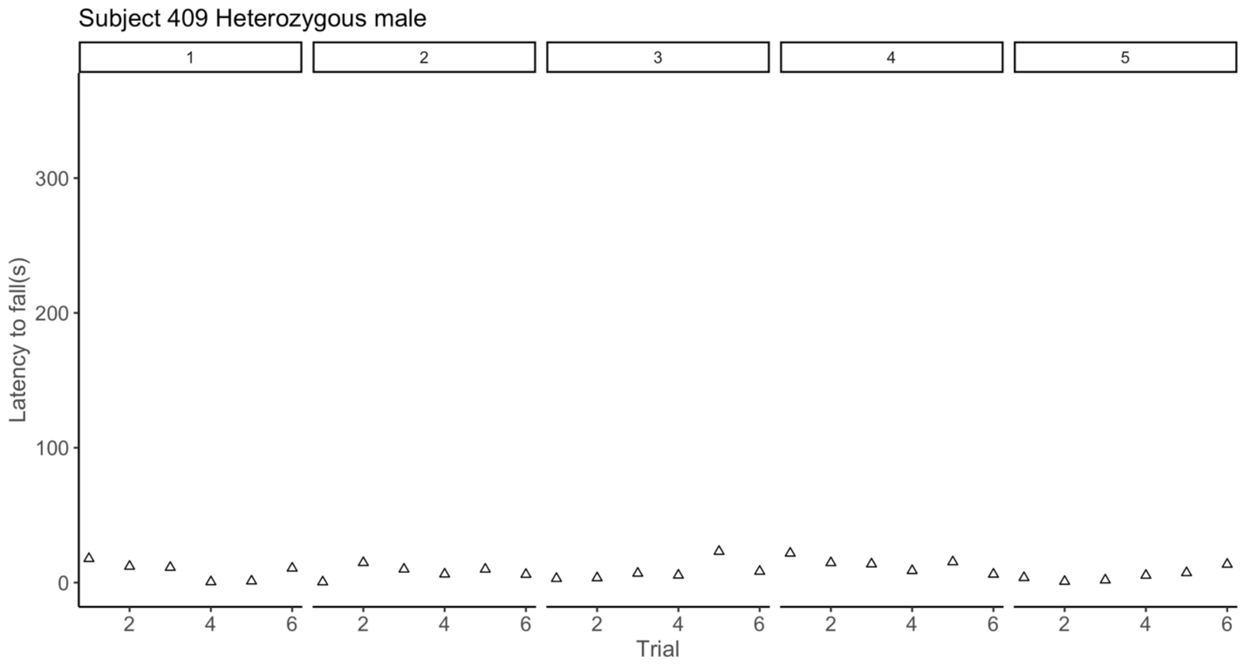


**S4 Fig. Latencies to Fall from the Rotarod of the Best and Worst Subjects.** Latencies (s) to fall from the Rotarod across days and trials for A) the best performing rat, a heterozygous female, and B) the worst performing rat, a heterozygous male.
