## Supplementary material for "Motor deficits in the McGill-R-Thy1-APP transgenic rat model of Alzheimer’s Disease": S4 Table

**S4 Table. Model selection table for distance travelled in the Y maze.** Akaike's Information Criterion (AIC) model comparison of distance travelled in the Y maze. Models are ranked from lowest to highest AIC values, and the degrees of freedom (df) used by the model calculated. The delta value indicates the difference between the AIC of a model and the AIC of the model with the lowest AIC. The weight of the models indicates the probability of the particular model out of all tested models.

| geno | sex | geno:sex | df | logLik | AIC | delta | weight |
|---|---|---|---|---|---|---|---|
| + | + | | 5 | -121.04 | 252.07 | 0.00 | 0.447 |
| | + | | 3 | -123.13 | 252.26 | 0.18 | 0.407 |
| + | + | + | 7 | -120.21 | 254.43 | 2.36 | 0.137 |
| | | | 2 | -128.18 | 260.35 | 8.28 | 0.007 |
| + | | | 4 | -127.72 | 263.43 | 11.36 | 0.002 |
