## Supplementary material for "Motor deficits in the McGill-R-Thy1-APP transgenic rat model of Alzheimer’s Disease": S5 Table

**S5 Table. Model Selection Table for Rat Weight**. Akaike's Information Criterion (AIC) model comparison of mean weight. Models are ranked from lowest to highest AIC values, and the degrees of freedom (df) used by the model calculated. The ∆ AIC value indicates the difference between the AIC of a model and the AIC of the model with the lowest AIC. The Evidence Ratio (ER) is calculated from the ∆ AIC values and indicates how much more likely the model with the lowest AIC is than each particular model. The weight of the models ($w_i$) is calculated from the evidence ratios and indicates the probability of the particular model out of all tested models.

| Model | *df* | AIC | Δ AIC | ER | $w_i$ |
|---|---|---|---|---|---|
| **Weight ~ sex** | 3 | 372.39 | 0 | 1 | 7.53e-1 |
| **Weight ~ sex + genotype** | 5 | 374.93 | 2.54 | 3.56 | 2.12e-1 |
| **Weight ~ sex × genotype** | 7 | 378.52 | 6.13 | 21.49 | 3.51e-2 |
| **Weight ~ 1** | 2 | 442.83 | 70.44 | 1.98e15 | 3.81e-16 |
| **Weight ~ genotype** | 4 | 445.28 | 72.90 | 6.74e15 | 1.12e-16 |
