## Supplementary material for "Motor deficits in the McGill-R-Thy1-APP transgenic rat model of Alzheimer’s Disease": S6 Table

**S6 Table. Model Selection Table for Maximum Latency to Fall from the Rotarod.** Akaike's Information Criterion (AIC) model comparison of maximum latency to fall off the Rotarod. Models are ranked from lowest to highest AIC values, and the degrees of freedom (df) used by the model calculated. The Δ AIC value indicates the difference between the AIC of a model and the AIC of the model with the lowest AIC. The Evidence Ratio (ER) is calculated from the Δ AIC values and indicates how much more likely the model with the lowest AIC is than each particular model. The weight of the models ($w_i$) is calculated from the evidence ratios and indicates the probability of the particular model out of all tested models.

| Model | *df* | AIC | Δ AIC | ER | $w_i$ |
|---|---|---|---|---|---|
| **Max Latency ~ weight + sex + genotype** | 6 | 378.13 | 0 | 1 | 2.53e-1 |
| **Max Latency ~ weight × sex + genotype** | 7 | 379.15 | 1.02 | 1.67 | 1.52e-1 |
| **Max Latency ~ weight + sex** | 4 | 379.70 | 1.56 | 2.19 | 1.26e-1 |
| **Max Latency ~ weight × genotype + sex** | 8 | 380.00 | 1.87 | 2.54 | 9.94e-2 |
| **Max Latency ~ weight × sex** | 5 | 380.54 | 2.41 | 3.34 | 7.57e-2 |
| **Max Latency ~ weight + genotype × sex** | 8 | 381.00 | 2.87 | 4.20 | 6.01e-2 |
| **Max Latency ~ weight** | 3 | 381.05 | 2.92 | 4.30 | 5.88e-2 |
| **Max Latency ~ (weight + sex + genotype)$^2$ - sex:genotype** | 9 | 381.17 | 3.04 | 4.57 | 5.53e-2 |
| **Max Latency ~ weight + genotype** | 5 | 381.91 | 3.78 | 6.63 | 3.81e-2 |
| **Max Latency ~ (weight + sex + genotype)$^2$ - weight:genotype** | 9 | 382.06 | 3.92 | 7.12 | 3.55e-2 |
| **Max Latency ~ (weight + sex + genotype)$^2$ - weight:sex** | 10 | 382.67 | 4.45 | 9.69 | 2.61e-2 |
| **Max Latency ~ weight × genotype** | 7 | 383.92 | 5.79 | 18.12 | 1.39e-2 |
| **Max Latency ~ (weight + sex + genotype)$^2$** | 11 | 384.21 | 6.08 | 20.93 | 1.21e-2 |
| **Max Latency ~ weight × sex × genotype** | 13 | 387.51 | 9.38 | 108.67 | 2.33e-3 |
| **Max Latency ~ sex** | 3 | 338.25 | 10.12 | 157.45 | 1.61e-3 |
| **Max Latency ~ genotype + sex** | 5 | 390.07 | 11.94 | 391.02 | 6.46e-4 |
| **Max Latency ~ genotype × sex** | 7 | 393.55 | 15.42 | 2234.29 | 1.13e-4 |
| **Max Latency ~ 1** | 2 | 396.25 | 18.12 | 8595.05 | 2.94e-5 |
| **Max Latency ~ genotype** | 4 | 396.71 | 18.58 | 10854.12 | 2.33e-5 |
